# Disruption of prefrontal cortex accelerates sensory anchoring but dissociates online recognition from implicit consolidation of sound sequences

**DOI:** 10.64898/2026.06.11.731781

**Authors:** Kurt D Shulver, Nicholas A Badcock, Paul F Sowman, David McAlpine, Heivet Hernández-Pérez

## Abstract

The auditory brain excels at extracting predictive regularities from sound, but the neural and temporal mechanisms mediating long-term learning of these regularities remain poorly understood. Here, using a perceptual anchoring paradigm, we investigate whether the dorsolateral prefrontal cortex (dlPFC) is causally required to transform transient regularities into stable internal templates (anchors). We characterized the behavioural dynamics of anchoring to predicable acoustic events using sensitivity (*d’*) and decision bias (*c*) metrics, alongside growth curve analysis (GCA) to model trial-by-trial learning. Listeners exhibited a robust anchoring effect, showing significantly higher sensitivity in decision strategy for recurring acoustic regularities compared to novel ones. This implicit anchoring was driven selectively by specific features in regular sequences—the local repetition of an acoustic motif—whereas explicit memory indiscriminately encoded the entire acoustic episode. To test for causality, we applied inhibitory continuous theta-burst stimulation (cTBS) to the dlPFC. Crucially, not only did this not impair anchoring, but significantly accelerated the rate of anchoring for predictable patterns, suggesting the dlPFC may act as a source of top-down interference in implicit learning. In contrast to predictable sequences, inhibition of dlPFC disrupted the initial precision of online comparisons for novel sequences, though insulating against the rapid, interference-driven reduction in performance or novel sequences in observed in listeners subjected to a sham cTBS procedure. These findings reveal a functional dissociation. Whilst dlPFC provides an essential buffer for novel sensory comparisons, long-term consolidation of regularities in acoustic sequences relies on a robust implicit pathway independent of prefrontal control.

## Introduction

Learning is critical to survival. We are constantly exposed to multiple streams of sensory information and must learn and act on those most relevant in a wide range of environments, situations, and contexts. Given metabolic and other constraints, it is likely that commonly and rarely occurring events are provided different perceptual weights, a phenomenon reflecting the ability of the nervous system to adapt to sensory information over multiple timescales. One means by which this is achieved is through statistical learning, an unconscious, automatic process by which regular sensory patterns are identified by virtue of the predictable nature of their appearance over time (Chun & Jiang, 1998; Frost et al., 2015; Kirkham et al., 2002; Saffran et al., 1996; Turk-Browne et al., 2009). Statistical learning (SL) appears to be a fundamental, domain-general principle of sensory organisation, enabling efficient processing across sensory modalities; vision (Fiser & Aslin, 2002; Turk-Browne et al., 2010), tactile sensation (Conway & Christiansen, 2005), and olfaction (Marcout et al., 2025; Tong et al., 2014), and in the sense of hearing (Agus et al., 2010; Saffran et al., 1996, 1999) where it is supports language acquisition (Aslin et al., 1998), the representation of background sounds ‘textures’ in terms of their summary statistics (Bastug et al., 2026; McDermott & Simoncelli, 2011; McWalter & McDermott, 2018), and in the parsing of complex room acoustics to better understand speech in background noise (Brandewie & Zahorik, 2010, 2013; Hernández-Pérez et al., 2025).

Whilst dedicated neural pathways to the level of auditory cortex are essential for processing the dynamic spectro-temporal features of sound (Banai et al., 2009; Bizley & Cohen, 2013; Souffi et al., 2021), higher-order statistical inference likely requires a distributed network involving corticofugal efferent pathways (Bajo et al., 2019; Souffi et al., 2021; Wang et al., 2023) as well as bidirectional communication with prefrontal brain regions that support the formation and maintenance of working memory, cognitive control, and multi-sensory integration (Ahveninen et al., 2023; Ambrus et al., 2020; Bonetti et al., 2024; Demarchi et al., 2019; Jang & Choi, 2022; Kumar et al., 2016; Plakke & Romanski, 2014; Rolls et al., 2023; Schiavo & Froemke, 2019; Winer & Lee, 2007). Nevertheless, despite clear anatomical evidence of bidirectional innervation of auditory and prefrontal cortical areas (Bidelman & Myers, 2020; Hockley & Malmierca, 2024; Petrides & Pandya, 2002; Zikopoulos & Barbas, 2006) such as dorsal-lateral prefrontal cortex (dlPFC), any causal role for these pathways in encoding long-term auditory regularities has not been established. Recently, we explored how the process of rapid learning of room acoustics improves understanding of speech in noise (Brandewie & Zahorik, 2010, 2013), and found this learning to be impaired when activity in dlPFC was disrupted (inhibited) with repetitive transcranial magnetic stimulation (rTMS; Hernández-Pérez et al., 2025). Importantly, listeners were not tasked with reporting any aspect of the abstract feature being learned—the reverberant characteristics of rooms—but were tasked with recalling key words from sentences spoken in those environments. Reverberation was characterised from the room impulse response (RIR), specifically the RT_60_, the time it takes for reflected sound energy to decay by 60 decibels. Controlled for semantic content, length of phrase, and talker identity, word recall improved with increasing exposure (over time) to a room, and this recall was impaired by continuous theta burst stimulation (cTBS) over dlPFC. We interpreted these findings as representing the learning of summary statistics, but in which a ‘bait and switch’ paradigm—speech understanding, rather than discrimination of the abstract statistical features being implicitly learned—demonstrates real-world benefit of implicitly learning a background feature defined by its statistics (reverberation time).

Here, we explore the contribution of dlPFC to statistical learning of abstract acoustic patterns in the framework of perceptual anchoring (Ahissar, 2007; Ahissar et al., 2006). Perceptual anchoring models learning as a hierarchical process in which short-term brain mechanisms track the statistics of unfolding sensory input (i.e., in the order of tens of milliseconds) to facilitate real-time identification, with longer-term mechanisms integrating this statistical information to generate a stable internal representation (perceptual anchor). In the context of listening, anchoring provides for more efficient recognition of sounds at a later time (Ahissar, 2007; Ahissar et al., 2006; Daikhin et al., 2017; Jaffe-Dax et al., 2018; Lieder et al., 2019). To do so, we employed a sound paradigm designed to minimise listeners’ abilities to encode explicitly regular features in an unfolding acoustic stimulus. The stimulus consists of rapid sequences of 50-ms pure tones—drawn from a limited pool of 20 frequencies—that transition from random (RAND) to regular (REG) segments at a fixed time point (Barascud et al., 2016). Whilst each RAND segment samples this pool uniformly, REG segments do so deterministically, in that they comprise a concatenated, local repeat (see Fig 1A). Transitions from RAND to REG segments provide a means of indexing the brain’s ability to process structural order at multiple temporal scales.

**Figure 1.**
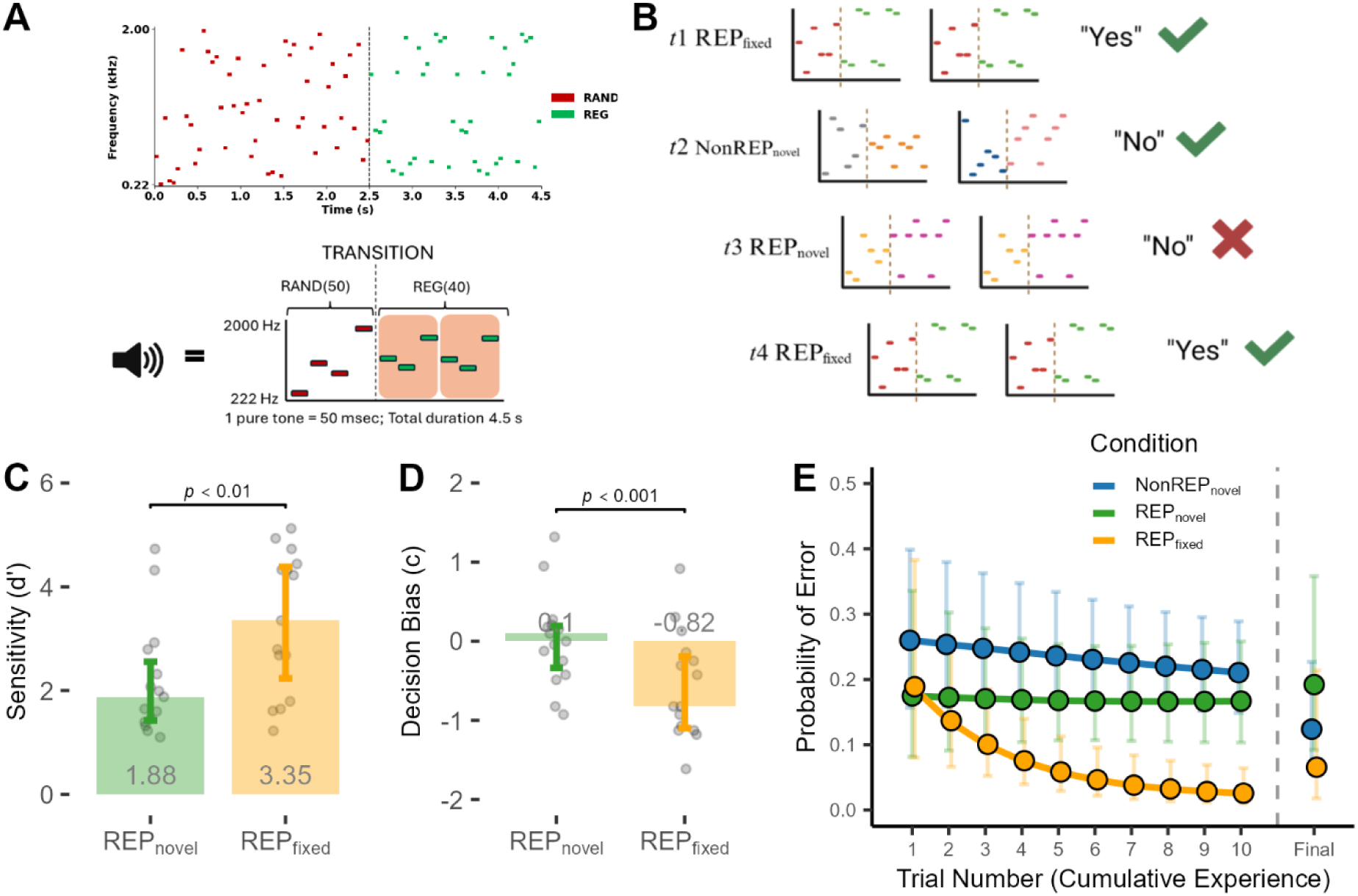
**(A: Top Panel)** *Sequence structure and composition.* Sequences (4.5 s total duration) comprised a random (RAND) segment of 50 stochastic tone-pips (2.5 s) followed by a regular (REG) segment (2 s). The REG segment consisted of two identical 20-pip segments presented without intervening silence. The repeition within the REG segment occurs at 3.5s, or 1 second into the REG segment. **(A: Bottom Panel)** *Condensed Representation of sequence structure and composition.* **(B)** Example of first 4 trials presenting the composition of sequences for each condition (REP_fixed_, REP_novel_, and NonREP_novel_) and correct responses for each. Participants were asked, “were these sounds the same?” As show, the correct response for REP sequences is “yes”, and the correct response for NonREP sequences is “no.” **(C)** IQR and individual data points showing overall sensitivity (*d’*). Participants were more sensitive to REP_fixed_ (med. =3.35, IQR [1.95, 4.35]) relative to REP_novel_ (med. = 1.88, IQR [1.44, 2.44]) **(D)** IQR and individual data points for overall criterion (*c*). REP_fixed_ condition, listeners adopted a significantly lower criterion (med. = −0.82, IQR [−1.10, −0.19]) relative to REP_novel_ (med. = 0.000, IQR [−0.34, 0.19]). A small, shaded area was added here to help visualise the median of 0. **(E)** *Trial-by-trial probability of FAs.* Solid lines represent fit of GCA model using 2^nd^-order orthogonal polynomials to capture non-linear trajectory of learning. Data points depict estimated marginal means at sampled intervals (Trials 1-10); vertical error bars indicate 95% confidence intervals. ‘Final’ trial number indicates estimated marginal means upon completion of the task.

Confirming that detection of transitions improves with familiarity, i.e., improved reaction times with increased exposure to the transition from RAND to REG (Barascud et al., 2016; Bianco et al., 2020), we found that REG, but not RAND, segments drove a sensory anchoring process. Listeners were increasingly able to discriminate whether repeated RAND+REG sequences in a trial were identical or not when these sequences were fixed (i.e. appeared repeatedly over the course of the experiment) compared to trials in which paired sequences of RAND+REG appeared only once (REP_novel_). Then, employing stimulus conditions in which the RAND (REP_RAND-fixed_) or the REG (REP_REG-fixed_) segments only of sequences were fixed across the experiment, we demonstrated that the anchoring effect was driven by the REG segment of our pure-tone sequences; i.e., anchoring relied on the locally in-time repeated ‘motif’ that constituted REG segments. Such was the ‘lure’ of REG segments to anchoring, that listeners mistakenly reported REP_REG-fixed_ sequences to be identical when their RAND segments were different; the reverse was not the case for the REP_RAND-fixed_ condition for which listeners accurately reported sequences in a pair to be different. This was despite listeners judging the same RAND and REG segments as being equally familiar in an explicit recognition task. Finally, hypothesising that integrating tone sequences into an anchoring framework is mediated by prefrontal cortical circuits, we applied continuous theta-burst stimulation (cTBS)—a rapid form of transcranial magnetic stimulation—to inhibit activity in dlPFC and determine any causal contribution of this brain region to perceptual anchoring. We found that disrupting dlPFC with cTBS speeds up the rate at which listeners accumulate knowledge of predictable auditory sequences to generate perceptual anchors but renders them less proficient at processing unpredictable sequences. Together, our data suggest that committing acoustic features to memory (anchoring) requires some form of regularity in their structure—local repetitions (REG segments) in acoustic structure that recur repeatedly—and that prefrontal cortical circuits modulate the rate and extent to which predictable and unpredictable sound sequences are committed to implicit memory.

## Methods

### Equipment

Perceptual anchoring tasks were programmed in MATLAB (ver. R2018b) using the Psychophysics Toolbox (Brainard, 1997; Kleiner et al., 2007; Pelli, 1997) and completed on a 15’ Dell laptop. Participants were seated comfortably in front of the laptop and headphones were inserted by the researcher. Sounds were presented over insert headphones (ER-2 insert phones, Etymotic) to both ears simultaneously. Stimuli were played through an RME Fireface UCX digital-to-analog converted with 16-bit resolution at a 44.1 kHz sample rate. All experiments were completed in an acoustically treated booth. The task was completed via key presses on the laptop keyboard.

### Acoustic Stimuli

Pure tone sequences were created as described by Barascud et al. (2016) and were generated using MATLAB (Mathworks ver. R2018b). Sequences were generated as segments of 50 ms tone-pips with frequencies drawn from a pool of 20 values equally spaced on a logarithmic scale between 222 and 2000 Hz. Each sequence consisted of two segments, a random (RAND) and regular (REG) segment. RAND segments consisted of 50 randomly generated tone-pips (2.5 s total duration). REG segments consisted of two recurring segments (20 tone pips each; 40 in total; 2 s total duration). The two recurring segments were identical in structure and content. There was no intervening silence between tone-pips and segments, making the total duration of sequences 4.5 s each. The experiment consisted of two conditions, a random and repeating condition. The random condition consisted of two NonREP_novel_ sequences that consisted of a novel RAND and REG segment in each sequence. This condition was nonidentical and non-recurring across the task. The repeating condition consisted of identical REP_novel_ or REP_fixed_ sequences. Both repeating conditions consisted of two sequences, however, REP_fixed_ sequences repeated across trials such that participants would hear the same sequence pair multiple times.

### Continuous Theta-Burst Stimulation

Participants were screened for contraindications to repetitive Transcranial Magnetic Stimulation (rTMS) prior to testing. Two continuous Theta Burst Stimulation (cTBS) conditions (“Active” and “Sham” TMS) were counterbalanced across participants, who were blinded to the type of manipulation they might receive.

CTBS was administer using a Magstim Rapid2 system (Magstim, Whitland, United Kingdom) and a 70-mm figure-of-eight coil oriented at 45° to the scalp with current flowing posterior-anterior across the primary motor cortex. cTBS intensity was individuals determined for each participant as a function of corticospinal excitability, assessed via single-pulse TMS motor thresholds. For threshold determination, we used visual observation of first dorsal interosseus (FDI) muscle twitches (Varnava et al., 2011). The coil position and angle were adjusted until the optimal site for consistent and robust FDI muscle twitch elicitation was identified. Participants were instructed to relax their arm on their lap, palm facing upwards, whilst gently squeezing their thumb and forefinger together. The minimum stimulator output producing a visible twitch in either the left or right hand was then determined, with a criterion of 3 out of 5 successful twitches in response to single-pulse TMS. Participants were occasionally asked to shake their arm to relax their muscles before resuming the hand position (Coltheart et al., 2018; Dienes & Hutton, 2013).

Motor thresholds ranged from 46-57% of maximum stimulator output. These individually determined thresholds were then used to set stimulation intensity for bilateral cTBS, delivered by placing the figure-of-eight-coil over the right and left dlPFC, located over electrode positions F3 and F4 of the 10-20 EEG system (Herwig et al., 2003; Jurcak et al., 2007). Bilateral stimulation was performed serially = targeting first either the right or left dlPFC (order counterbalanced across participants) – to limit compensatory activity from the non-stimulated hemisphere (Ambrus et al., 2020). Each cTBS burst consisted of three pulses at 50 Hz, with bursts repeated at 5 Hz and applied continuously for 40 s at an intensity of 80% of the individual resting motor threshold (Huang et al., 2005). This protocol elicits a period of depressed cortical excitability lasting approximately one hour post-stimulation (Gamboa et al., 2010; Hoogendam et al., 2010), the window during which anchoring to pure-tone sequences was assessed. Our overall neurostimulation setup and technique are consistent with previous from your group (Hernández-Pérez et al., 2025)

### Procedure

The experiment consisted of a 2-alternative-forced-choice (2AFC) task and contained two phases: a training phase followed by a test phase. Training was completed to allow participants to familiarise themselves with the task and to be better able to distinguish NonREP and REP conditions. For each trial within a phase, participants were exposed to two stimuli separated by an interstimulus interval (ISI; 2.0 s). Participants were asked “were the sounds the same?” (i.e., were the two sequences identical; ‘Yes’ for they were the same, and ‘No’ for they were not the same). Responding with a key press (“f” for Yes, “j” for No) triggered the next trial until all trials were completed. Participants were provided a single break which was predetermined (i.e., occurred half-way through test phase) and self-directed (i.e., lasted as long as the participant desired). Training phases included only NonREP_novel_ and REP_novel_ conditions. The test phase followed an identical procedure to the training phase but included the additional REP_fixed_ condition. Significantly, participants were not instructed that the REP condition contained a reoccurring stimulus pair (REP_fixed_), only that they should report when they perceived the two pure-tone sequences as the same.

### Statistical analysis

Behavioural data were analysed using Linear Mixed-Effects Models (LMMs) and Generalized Linear Mixed-Effects Models (GLMMs). Mixed-effects modelling provides a robust alternative to traditional repeated-measures ANOVA by incorporating both fixed-effects (e.g., Condition, Group) and random-effects terms. This approach allows for the statistical description of the sample population while simultaneously quantifying inter-participant and inter-item variability. Mixed models are particularly suited for this paradigm as they effectively accommodate unbalanced designs and account for the non-independence of data points nested within individual participants (Magezi, 2015).

All data analysis and figure generation were completed though a combination of packages in R (R Core Team, 2019): Tidyverse for data wrangling and visualisation (Wickham & RStudio, 2023); lme4 and lmerTest for mixed effects modelling (Bates et al., 2015; Kuznetsova et al., 2017); and emmeans for post-hoc comparisons and estimated marginal means (Lenth et al., 2026; Searle et al., 1980). For specific non-linear estimation and convergence analysis, we utilised nlstools (Baty et al., 2015), nlsmultistart (Padfield et al., 2025), and nlsr (Nash et al., 2026). Full model output for all analyses, including fixed and random effects estimates, standard errors, and degrees of freedom, is reported in the Supplemental Information.

The primary outcome measure for task proficiency was d-prime (*d’*), a sensitivity index derived from signal detection theory (Macmillan & Creelman, 2004; Stanislaw & Todorov, 1999). Sensitivity for each condition was calculated based on hit rates (*H*) for REPnovel and REPfixed trials, a shared false-alarm rate (*FA*) derived from the NonREPnovel foil condition, and a function for generating a normal distribution (*z*; *M* = 0, *SD* = 1):

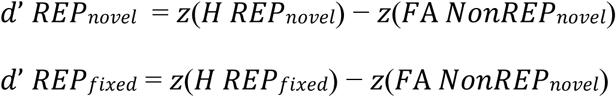

As REP_novel_ and REP_fixed_ stimuli were randomly interspersed within the same experimental block, they share a common NonREP_novel_ baseline. For cases where hit or false-alarm rates reached 0 or 1, a standard correction of +-0.1 trials was applied to allow for z-score calculation.

### Growth Curve Analysis (GCA)

To evaluate the temporal dynamics of learning, we employed a Logistic Growth Curve Analysis (GCA). Trial-by-trial error probabilities were modelled using a binomial link function (logit). The temporal trajectory of learning was captured using orthogonal polynomials, allowing for the independent evaluation of the overall reduction in error rate (i.e., improvement) (linear time term) and the stabilisation or “plateauing” of performance (quadratic time term). For these models, we checked whether the large estimates were caused by participants reaching a performance floor. When a group makes almost no errors, the model’s math can become unstable, producing inflated numbers that reflect this lack of variance rather than a literal rate of change. In such cases, model estimates were grounded in raw descriptive statistics and participant-level distributions to ensure interpretive rigor.

## Results

Successful sensory anchoring reflects a combined process of adaptation by a perceptual system to stimulus-relevant information and recruitment of attentional mechanisms to select, separate, and store that information for later access (Ahissar, 2007; Ahissar et al., 2006; Lieder et al., 2019). To assess the capacity of listeners to form perceptual anchors we employed sequences of pure tones comprised of distinct random (RAND) and regular (REG) segments. RAND segments were constructed from 50 randomly generated tone-pips (2.5-s total duration), and REG segments from two consecutive and identical motifs of 20 tone-pips (40 in total; 2-s total duration) (Figure 1A). No intervening silence separated RAND and REG segments in a sequence, or the motifs in the REG segment, providing for a total sequence duration of 4.5 s. Each trial consisted of two complete sequences, separated by an interstimulus interval (ISI) of 2.0 s. In a REP_fixed_ condition, pairs of sequences in a trial were identical (including RAND and REG segments) and appeared multiple times (i.e., were ‘fixed’) across the course of the experiment. In a REP_novel_ condition, pairs of sequences in a trial were also identical (i.e., repeated) but appeared only once during the experiment (’novel’). In a third condition, NonREP_novel_, motifs within each REG segment of a sequence were identical, but both the RAND and the REG segments differed between the two sequences presented in a trial, and these sequences never reappeared during the experiment.

### Listeners anchor to repeated, fixed sequences of pure tones

We first established whether listeners could utilise the recurrence of pure-tone sequences to improve their recognition of these sequences, even in the absence of explicit instruction to do so. In a two-alternative forced choice (2AFC) task, 16 normal-hearing (M_age_, = 33.1 SD = 5.16) participants were exposed to an experimental block of 80 trials. For REP_fixed_ and REP_novel_ conditions the correct response was ‘yes’ (the sequences were the same), and for NonREP_novel_ the correct answer was ‘no’ (the sequences were different). Prior to testing, all participants completed a brief training session to familiarise them with the task. Importantly, REP_fixed_ stimuli were not presented to listeners during the training phase (data obtained during the training phase are included in the Supplemental Information).

Performance in the test phase of the task was first analysed in terms of global sensitivity (*d’*; d-prime) and decision bias (*c*; criterion), focusing on the magnitude of the anchoring effect between REP_fixed_ and REP_novel_ conditions. Generalised linear mixed-effects modelling (GLMM) was used to quantify these fixed effects of condition whilst accounting for random effects associated with individual listeners (Bates et al., 2015; Kuznetsova et al., 2017). This approach allowed us to estimate the variability participants contributed to overall performance, with Cohen’s d values reported to specify effect sizes (Cohen, 1988, 2013). Because global false-alarm (FA) rates (derived from the NonREP_novel_ condition) served as a shared baseline for REP_fixed_ and REP_novel_ conditions, variance in both sensitivity and decision bias was driven entirely by fluctuations in hit rates of those specific conditions. Consequently, shifts in decision bias are mathematically proportional to shifts in sensitivity within our models.

Our data confirmed a robust anchoring effect (Figure 1C): listeners were significantly better at detecting whether the two sequences comprising a trial were identical when these paired sequences appeared consistently across an experimental block (REP_fixed_; *d’* = 3.35) compared to when an identical pair of sequences appeared only once (REP_novel_; *d’* = 1.87). This improvement in performance was substantial (*p* < .001), with a large effect size (d = 1.02), confirming that listeners were able to exploit long-term regularities in sequences of pure tones to identify paired sequences as being identical (see Tables S1–S3, Supplemental Information).

Improvements in recognising repeated spectro-temporal features of pure-tone sequences with increasing exposure might signify a change in perceptual acuity but they can also represent a shift in decision strategy—the internal threshold, or criterion (*c*), used to categorise a stimulus. We therefore analysed decision bias (*c*) to determine how listeners’ willingness to report a repeated sequence as actually being repeated evolved as the anchor was formed (Figure 1D). For the REP_novel_ condition—in which tone sequences were never experienced previously—listeners remained essentially unbiased (*c* = 0.00), suggesting an equal weighting of decisions for each unique sequence. However, in the REP_fixed_ condition, the formation of an auditory anchor fundamentally altered their approach to the task, with listeners adopting a significantly more liberal criterion (*c* = −0.82, *p* < .001), indicating a shift in confidence where they became primed to say ‘yes’ to the learned pattern.

To capture the evolution of performance in the task as a function of time (i.e., the rate of learning), we employed a logistic growth curve analysis (GCA) to model trial-by-trial error probabilities. We focused on error rates rather than aggregate *d’* to isolate the temporal dynamics of how listeners refine their internal representation for fixed vs novel sequences. This approach allows us to assess learning trajectories by independently evaluating the overall rate of improvement (linear time component) and the stabilisation of performance (quadratic time component) as a function of time spent within the condition (i.e., cumulative exposure). This analysis revealed fundamentally different learning trajectories for the different sound structures (Figure 1E). For REP_fixed_ sequences, listeners exhibited a rapid ‘locking-in’ phase, characterised by a sharp reduction in errors (i.e. reporting sequences in a pair to be different when they were in fact identical) during initial exposures (approximately trials 1-10) followed by a stable plateau (*p* < .05). This non-linear trajectory reflects the swift consolidation of a stable auditory anchor. In contrast, the REP_novel_ condition followed a flat linear trajectory (*p* =.40); in the absence of a recurring pattern to accumulate, listeners had to rely on real-time processing of acoustic information to make their decision for each trial individually. Error rates in the NonREP_novel_ condition (false alarms) showed a slow, gradual decline, suggestive of a general task-learning effect whereby listeners refine their criteria for what constitutes a ‘different’ sequence.

### Perceptual anchoring is driven by locally repeated motifs within fixed sequences

Listeners appear rapidly to ‘lock in’ to auditory patterns that re-appear over time, rendering features within these patterns more easily discriminable. Specifically, spectro-temporal sequences are more likely to be judged as doing so if they appear over the course of an experiment—i.e., they become anchored. However, it remains unclear which segments, or features of those segments, within a sequence supports the anchoring process. Thus far, our sequences comprise two distinct segments: RAND, consisting of a non-recurring pattern of tones, and REG, containing a locally recurring pattern of tones—i.e. two concatenated instances of a ‘motif’ (Figure 1A). REG segments—which always comprise the second half of a sequence—are distinguishable from RAND segments by this locally repeating motif. Nevertheless, RAND segments in REP_fixed_ sequences are identical within and across trials and could provide for perceptual anchoring if listeners were able to judge this to be so and, importantly, exploit this knowledge to form a perceptual anchor. To determine whether listeners adopt this strategy, we explored whether anchoring relies on global sensitivity to the entire acoustic sequence, including the RAND segment, or whether anchoring is dominated by, or exclusively the preserve of, the REG segment. To do so, we introduced two additional conditions to our standard paradigm (Figure 2A). In a REP_REG-fixed_ condition the REG segment remained identical between sequences in a trial and across trials, but the RAND segment differed between sequences in a trial; conversely, in the REP_RAND-fixed_ condition, the RAND segment was identical between sequences in a trial and fixed across trials, and the REG sequence differed. Importantly, the respective RAND and REG segments were identical to those of the REP_fixed_ condition, thereby doubling the opportunity for listeners to become familiar with individual segments. This design allows for a specific behavioural prediction: if listeners successfully anchor to the REG segment in the REP_REG-fixed_, they may fail to detect the change in the varying RAND segment, resulting in an error (i.e., judging the sequences as identical when, by means of the different RAND segments between the two sequences, they are not). Thus, a drop in sensitivity (negative *d’*) in these conditions serves as an indicator of the strength of the entire fixed sequence, not only just the repeated structure of the REG segment, in driving anchoring.

**Figure 2.**
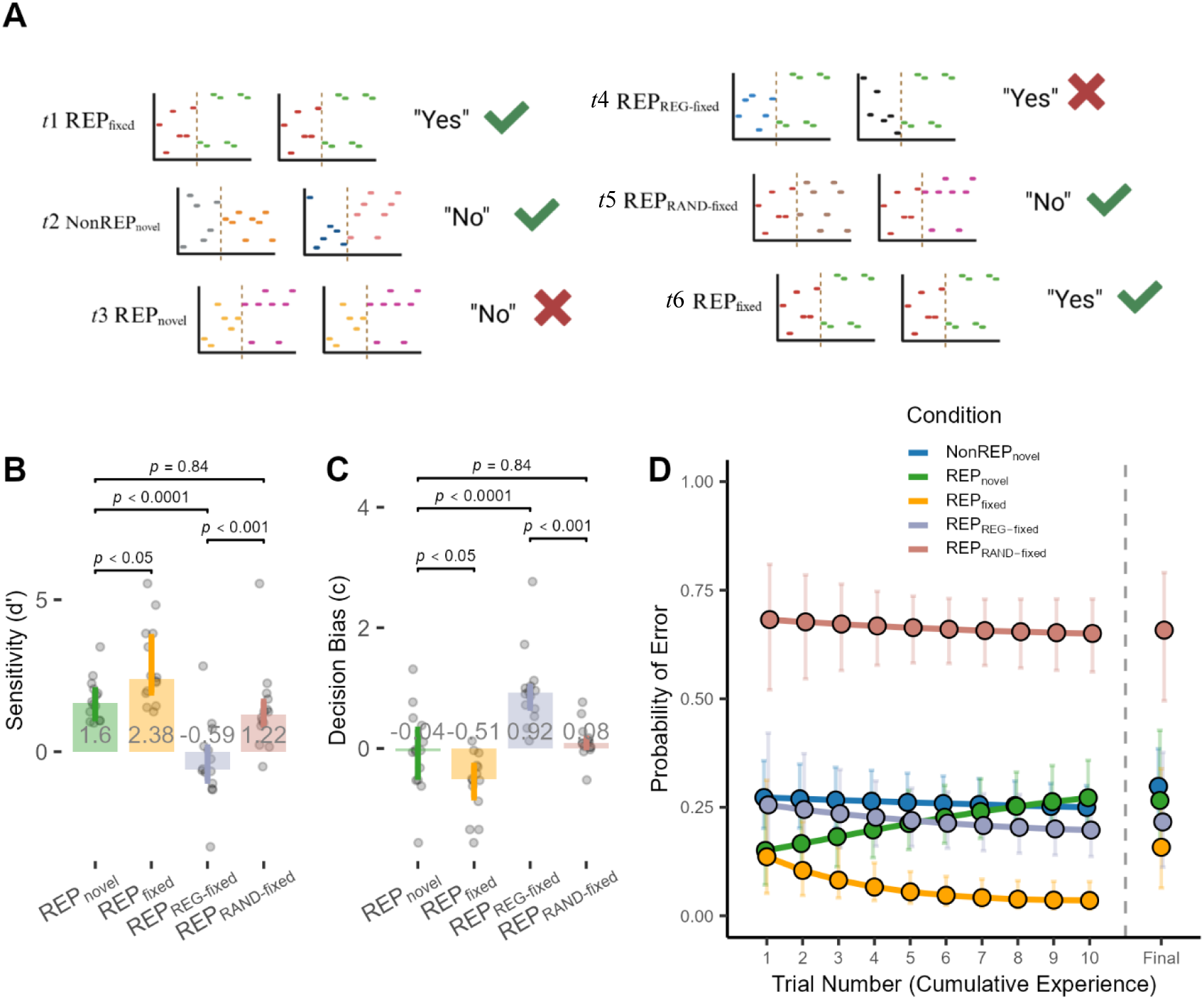
**(A)** *Procedure for sequence presentation within and across trials.* An example illustration of the first 6 trials presenting the composition of sequences for each condition (REP_fixed_, REP_novel_, and NonREP_novel,_ REP_REG-fixed,_ REP_RAND-fixed_) and the correct response for each condition. Participants were asked, “were these sounds the same?” REP_REG-fixed,_ and REP_RAND-fixed_ are made up of the REP_fixed REG_ and _RAND_ segments, but with novel _RAND_ and _REG_ segments, respectively. **(B)** *Figure for results depicting sensitivity (d’) per condition.* Bar plots include error bar representing the interquartile range, and jitter points showing overall sensitivity (*d’*) for each participant: REP_fixed_ (Med =2.38, IQR [1.95, 3.78]); REP_novel_ (Med = 1.59, IQR [1.10, 2.02]), REP_REG-fixed_ (Med = −0.59, IQR [−0.95, 0.14]), REP_RAND-fixed_ (Med = 1.22, IQR [0.87, 1.63]). **(C)** *Figure for results depicting criterion (c) per condition.* Bar plots include error bar representing the interquartile range, and jitter points showing overall criterion (*c*) for each participant: REP_fixed_ (Med = −0.51, IQR [−0.81, −0.29]); REP_novel_ (Med = −0.04, IQR [−0.47, 0.31]); REP_REG-fixed_ (Med = 0.92, IQR [0.67, 1.02]), REP_RAND-fixed_ (Med = 0.08, IQR [0.02, 0.16]). **(D)** *Trial-by-trial probability of false alarms across conditions.* The solid lines represent the fit of the GCA model using second-order orthogonal polynomials to capture the non-linear trajectory of learning. Points depict estimated marginal means at sampled intervals (Trials 1-10), with vertical error bars indicating 95% confidence intervals. The “Final” trial number indicates the estimated marginal means at the end of the task.

We confirmed the initial anchoring effect in a new cohort of listeners (*N* = 14; *M* = 20.4, *SD* =1.70), finding, as before, that performance was significantly enhanced for the REP_fixed_ (*d’* = 2.38, *p* < .05) compared to the REP_novel_ (*d’* = 1.60) condition (Figure 2B; see Tables S4–S6, Supplemental Information), and that the full REP_fixed_ sequence induced a significantly more liberal decision bias (*c* = −0.51) compared to the unbiased REP_novel_ baseline (*c* = −0.04; *p* < .05) (Figure 2C). We then explored which of the two segments in a sequence—RAND or REG—underpinned anchoring and observed a striking dissociation in the contribution of the two segments. Specifically, when the RAND segment was fixed but the REG segment differed (REP_RAND-fixed_), listeners generally correctly identified the two sequences in a trial as different (*d’* = 1.22); their overall sensitivity and decision bias (*c* = 0.08; *p* = .84) did not differ significantly from the completely novel baseline (i.e., REP_novel_). This suggests that the RAND segment alone was not sufficient to elicit anchoring. Conversely, when the REG segment was fixed but the RAND segment changed (REP_REG-fixed_), listeners exhibited a profound collapse in sensitivity (*d’* = −0.59), performing significantly worse at determining or not whether a pair of sequences was repeated than for the novel baseline (REP_novel_; *p* < .001) or the REP_RAND-fixed_ (*p* < .001) *conditions*. This reduction in sensitivity was accompanied by a pronounced conservative shift in decision bias for the REP_RAND-fixed_ *condition* (*c* = 0.92), which differed significantly from all other conditions (*p* < .01). The data are consistent with the interpretation that listeners could not reliably distinguish different RAND segments in a sequence when the REG segments were identical. Instead, the fixed REG segment acted as a powerful perceptual cue, listeners becoming so anchored to the REG segment that this exerted an overwhelming influence on their perceptual judgements. This confirms that the REG segment is the dominant feature driving the anchoring effect, even when the same RAND segment reappears over the course of the experimental block (i.e., in the REP_RAND-fixed_ and REP_fixed_ configurations).

To determine if the partial anchor sequences—REP_RAND-fixed_ and REP_REG-fixed_—shared the same rapid learning profile as the complete sequences, we once more applied logistic GCA to the trial-by-trial error rates (Figure 2D). The full REP_fixed_ condition showed a significant non-linear trajectory (*p* < .01) for this group of participants, characterised by a rapid reduction in errors followed by a period of stabilisation (trials 8-10). Again, the REP_novel_ and NonREP_novel_ conditions showed no evidence of this stabilisation; their trajectories lacked the significant positive deceleration observed for fixed patterns, reflecting the continuous, linear demand of processing trial-unique information. Crucially, like the REP_fixed_ condition, the REP_REG-fixed_ condition—in which the RAND segment differed between sequences in a trial—also exhibited a significant non-linear learning trajectory (*p* <.05), mirroring the stabilisation of the full sequence anchor. Conversely, however, the REP_RAND-fixed_ condition did not show a clear non-linear trajectory (*p* = .052), behaving more like the flat, linear profile of the REP_novel_ condition. This confirms that, absent the locally repeating motif of the REG segment, listeners seem unable to establish a stable temporal prediction of pure-tone sequences, even when the RAND segment is fixed (i.e., identical across sequences and trials).

Having established that anchoring is driven largely by the REG segment, we next sought to determine whether this structural learning translated to explicit, conscious memory. To assess this, listeners completed a brief 2AFC recognition task following the discrimination task. They were asked to judge whether trials contained full sequences (e.g., REP_fixed_) or isolated segments (REP_REG-fixed_) had been presented in an experimental block. Due to the limited number of trial-unique exposures in this phase, we restricted our analysis to descriptive group-level percentiles (condition-specific analysis is available in Supplemental Information). Specifically, we report both accuracy (the proportion of objectively correct judgements) and endorsement rate (the proportion of ‘yes’ responses, regardless of accuracy). Together, these metrics allows us to distinguish between genuinely explicit memory of the sequences and any underlying response bias in listeners’ judgements of whether they previously experienced a specific sequence or segment.

Notably, explicit recognition did not mirror the selective nature of the implicit anchoring effect. Listeners were highly accurate at explicitly recognising the full, intact REP_fixed_ sequences (91.7% accuracy), but their memory for isolated segments revealed an intriguing dissociation. Although the REG segment exclusively drove the behavioural advantage in experiments in which listeners experienced REP_fixed_, NonREP_novel_ and REP_novel_ only, listeners were equally likely to explicitly endorse the previously heard RAND segment from REP_RAND-fixed_ (83.3%) as they were the REG segment from REP_REG-fixed_ (83.3%). Further, when presented with completely novel segments (i.e., segments never presented in the prior task), listeners exhibited a high error rate (endorsing novel segments 66.7% of the time), reflecting a bias in decision-making and poor overall accuracy.

Together, our data demonstrate a clear dissociation between implicit statistical learning and explicit auditory memory. In the first (discrimination) task, the formation of a stable auditory anchor is driven principally by the REG segment, with its concatenated identical motifs rather than any global sensitivity to the entire sequence, including the RAND segment. When only the REG segment was fixed across trials (REP_REG-fixed_), listeners exhibited the signature rapid learning trajectory, and this segment was so perceptually dominant that it resulted in listeners ignoring or missing within and between trial changes in the RAND segment. In contrast, when the REG pattern varied, learning remained flat (i.e., linear) even when the RAND segment was fixed (REP_RAND-fixed_). This confirms that the function describing the learning process for REP_fixed_ in the initial paradigm (REP_fixed_, REP_novel_, NonREP_novel_) relies largely on the availability of the repeating (REG) segment. However, this structural precision is lost under conscious recall. Whilst the implicit system responsible for statistical learning selectively extracts the predictable segment to facilitate real-time discrimination, explicit memory indiscriminately encodes the broader acoustic episode, treating both predictable (REG) and unpredictable (RAND) patterns as equally familiar. Thus, without a stable deterministic pattern to drive implicit computation, the auditory system cannot establish a predictive anchor; even when random noise is repeated across trials (as in REP_RAND-fixed_), listeners are forced to process the sequences as fundamentally novel events, mirroring the baseline performance of the REP_novel_ condition.

### Inhibiting activity in dorsolateral prefrontal cortex disrupts processing of regularity within acoustic sequences

We have demonstrated that listeners rapidly ‘lock in’ to auditory regularities—specifically locally repeated motifs forming the REG segments of our tone sequences—an implicit anchoring effect driven principally by some aspect of their predictability (Figure 1A). To explore any causal neural architecture supporting this implicit form of learning, we applied inhibitory continuous theta burst stimulation (cTBS) bilaterally over dorso-lateral prefrontal cortex (dlPFC) prior to participants undertaking the anchoring task, with the aim of temporarily disrupting local cortical excitability. We applied bilateral cTBS to an ‘active’ group of listeners (*N* = 10) serially across the left and right dlPFC to prevent compensatory mechanisms from the non-stimulated hemisphere. A separate ‘sham’ control group (*N* = 10) received identical bilateral procedures using a matched sham ‘figure-8’ coil to the active coil (Magstim) that delivers < 5% of the active stimulator output. Crucially, both coils mechanically generate an identical audible ‘click’ during the pulse trains (three pulses at 50 Hz, repeated at 5 Hz), providing a means of blinding listeners to their stimulation condition. We then evaluated anchoring proficiency in this new cohort of listeners, comparing sensitivity (*d’*) between the REP_novel_ and REP_fixed_ conditions and contrasting performance between active and sham groups. Additionally, we applied GCA to trial-by-trial error rates and assessed the time-to-convergence to determine whether activity in dlPFC modulates the temporal dynamics of learning.

We first assessed whether disruption of dlPFC with cTBS impairs the overall proficiency of perceptual anchoring. It did not. Both the active and sham groups exhibited a robust anchoring effect, demonstrating significantly greater sensitivity to predictable REP_fixed_ sequences relative to the REP_novel_ baseline (Figure 3A). Specifically, the sham group showed a clear advantage in sensitivity for the predictable spectro-temporal patterns of the REP_fixed_ sequences (*d’* = 2.66) over the unique patterns of the REP_novel_ sequences (*d’* = 1.56; *p* < .01). Crucially, this advantage was preserved in the active TMS group, which showed an almost identically enhanced sensitivity for REP_fixed_ (*d’* = 2.64) compared to REP_novel_ (*d’* = 1.46; *p* < .01) sequences. The interaction between *condition* (REP_fixed_ *vs*. REP_novel_) and *group* (active *vs*. sham) was not significant. The significant and positive main effect of *condition* (*d* = 1.72) is consistent with the robust implicit anchoring advantage established we have observed, and the near-zero effect size for group (*d* = −0.001) confirms that overall signal detection was unperturbed following modulation of dlPFC with cTBS (see Tables S7–S13, Supplemental Information).

**Figure 3.**
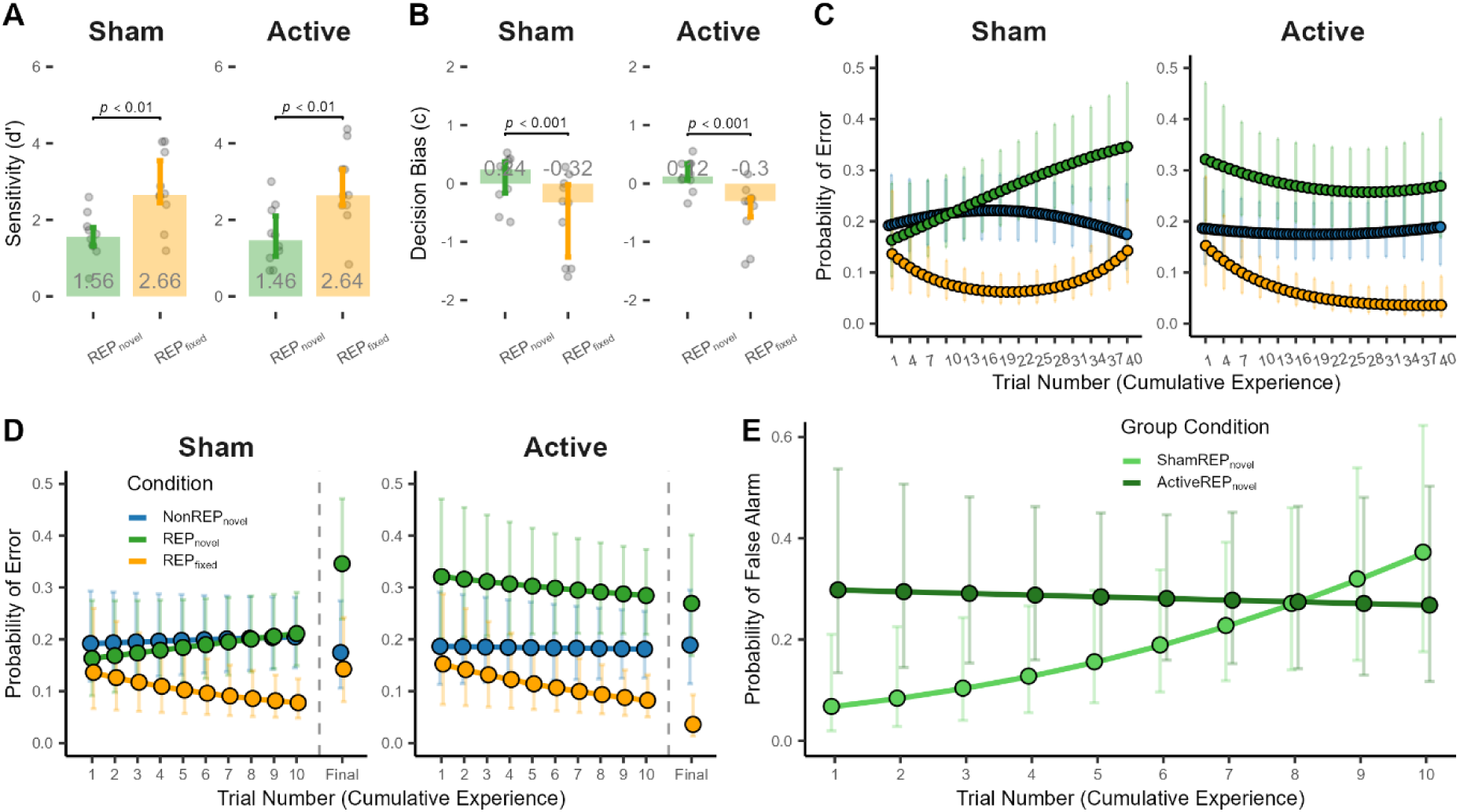
**(A)** *Figure for results depicting sensitivity (d’) per condition.* Bar plots include error bar representing the interquartile range, and jitter points showing overall sensitivity (*d’*) for each participant. Separated by group: Sham TMS and Active TMS **(B)** *Figure for results depicting criterion (c) per condition.* Bar plots include error bar representing the interquartile range, and jitter points showing overall criterion (*c*) for each participant: Separated by group. **(C)** *Trial-by-trial probability of false alarms across conditions and between Groups.* The solid lines represent the fit of the GCA model using second-order orthogonal polynomials to capture the non-linear trajectory of learning. Points depict estimated marginal means at sampled intervals (Trials 1-40), with vertical error bars indicating 95% confidence intervals. The 80 NonREP trials have been normalised to fit within the 40-trial axis length. **(D)** *Early trial probability of false alarms across conditions and between Groups.* **(E)** *Early trial probability of false alarms for REP_novel_ between Groups*.

As well as overall sensitivity in discrimination performance, the characteristic shift in decision bias also survived inhibition of dlPFC with cTBS, with listeners in the sham group adopting a significantly more liberal response bias for REP_fixed_ (*c* = −0.31) compared to REP_novel_ (*c* = 0.23) sequences (Figure 3B). Listeners in the active TMS group exhibited a similar strategic shift, moving from a baseline bias of 0.124 for REP_novel_ to a liberal bias of −0.32 for the REP_fixed_ anchor. However, whilst the sham group exhibited a wider interquartile range for decision bias in the REP_fixed_ condition, this variance was primarily driven by a ceiling effect, wherein a subset of listeners achieved perfect discrimination, pushing their criterion estimates to the bounds of the criterion level (see Supplemental Information).

We applied a logistic GCA to the trial-by-trial error rates to evaluate the impact of cTBS on long-term structural recognition of sequences. A significant triple interaction emerged between *group*, the REP_fixed_ condition, and the linear *time* term—representing the constant rate of error reduction across the experimental block (*b* = −94.60, *SE* = 42.47, *z* = −2.23, *p* < .05). Specifically, cTBS significantly accelerated the rate of anchoring for REP_fixed_ patterns relative in the active group relative to the sham group. Nevertheless, the extreme magnitude of this coefficient (*b* = −94.60) reflects the model’s attempt to map a near-total collapse in inter-participant variability as the cTBS group reached a performance ceiling. Specifically, the error (raw FA) rate for the cTBS group in the REP_fixed_ condition dropped from 0.14 (*SD* = 0.35) in the early phase of the task to 0.02 (*SD* = 0.14) by the end, with 60% of participants achieving a perfect zero-error rate. In contrast, the error rate and variance of the sham group remained unchanged (*M* = 0.14, *SD* = 0.35). Whilst aggregate sensitivity (*d’*) confirmed that participants in both groups eventually anchored to the fixed regularity (several sham participants achieved *d’*> 4.0), the GCA reveals that this anchoring was significantly more efficient when dlPFC was inhibited following cTBS. The *d’* scores, as a summary of the entire experimental block, likely obscure these differences due to the ceiling effects reached by both groups.

To investigate how inhibition of dlPFC modulated online recognition—i.e. REP_novel_ in which no trial-to-trial learning is possible—we used an additional GCA to model the first 10 trials (*trial)* of each condition (Figure 3D). This model revealed a significant three-way interaction between *group*, the REP_novel_ *condition*, and *trial* (*z* = −2.24, *p* < .05). Notably, no such interaction was observed for the REP_fixed_ condition during this 10-trial window (*p* = .93). This suggests a temporal dissociation in the effects of cTBS; whilst the advantage for REP_fixed_ sequences emerges through late-stage stabilisation, the modulation by TMS of the processing of REP_novel_ sequences is a transient effect restricted to early trials in the task. A subsequent *post-hoc* analysis of the REP_novel_ condition demonstrated that the between-group comparison of the rate of learning (slopes of task performance as a function of time) for this condition—the interaction between *group* and *trial* (Trials 1-10)—did not reach statistical significance (*p* = .52). However, an evaluation of within-group slopes—quantifying performance change over time for each group independently—revealed a clear divergence in stability of learning between the sham and active groups (Figure 3E). Specifically, the sham group exhibited a significant reduction in performance for the REP_novel_ condition; error probability rose steadily from 6.8% at Trial 1 to 37.2% by Trial 10 (*z* = 2.42, *p* < .05). In contrast, performance in the cTBS group, was overall higher to begin with, but remained stable throughout this early phase, with predicted error probability shifting only marginally from 29.8% to 26.8% (*z* = −1.99, *p >* .05, n.s.). Performance in the NonREP_novel_ condition was structurally similar across groups, suggesting this elevated error rate does not reflect a generalised deficit in auditory discrimination when dlPFC is disrupted. Rather, whilst cTBS initially disrupts the demanding online computation required for REP_novel_ comparisons, it simultaneously seems to insulate performance against the interference-driven decay observed in the sham group.

Together, the data reveal a functional dissociation in neural mechanisms contributing to the processing of auditory sequences. Inhibiting dlPFC not only preserves the ability of listeners to extract recurring regularities in acoustic sequences, but it also significantly accelerates the anchoring process, driving listeners to a performance ceiling faster than is observed when activity of dlPFC is intact. This suggests that whilst dlPFC is not necessary for implicit consolidation of fixed regularities, its typical activity may modulate processes that slow the transfer of fixed patterns to long-term memory. In contrast, when processing novel, trial-unique repetitions (REP_novel_), disruption of dlPFC leads to an initial deficit in accuracy with which novel acoustic patterns are detected but insulates against the rapid reduction in performance for REP_novel_ sequences observed in sham listeners. This suggests a clear trade-off in functionality when activity in dlPFC is intact compared to when it is not. Specifically, dlPFC may provide an online buffer required for precise novel comparisons, yet this same monitoring mechanism is prone to the interference-driven collapse observed as task demands accumulate (i.e., learning of REP_fixed_ sequences becomes complete). Ultimately, while dlPFC appears to be required for maintaining and comparing novel sensory input in real-time, the mastery of repeating auditory anchors operates via a robust, implicit pathway that is not only independent of prefrontal cortical control but is optimised in its absence.

## Discussion

We investigated the perceptual, cognitive, and neural architectures underlying the rapid perceptual anchoring to structured auditory sequences. We sought to determine if the auditory brain can extract complex regularities from sequences of acoustic patterns, the structural features contributing to this stabilisation, and whether activation of dorsolateral prefrontal cortex (dlPFC) is causally required for this process to occur.

We confirmed that listeners rapidly extract and ‘lock in’ to specific structural features, storing information about these features for future retrieval. Listeners increasingly judged paired sequences of pure tones as being identical if these sequences appeared multiple times across the course of an experiment (i.e., were ‘fixed’) compared to paired sequences that were also identical but appeared only once (’novel’) over the experiment. Isolating the specific driver of this effect, we revealed a striking behavioural dissociation: anchoring is not driven by global sensitivity to the entire acoustic sequence but, rather, by the perceptual dominance of the locally (in time) repeated motif of tones in the REG segment of REP_fixed_ sequences. Such was the influence of this perceptual cue that listeners frequently failed to notice trial-unique changes in the immediately preceding pattern (the RAND segment of the REP_REG-fixed_ sequence). Though specificity for the REG segment, but not the RAND segment, appeared to be implicit, a conscious recognition task revealed that both the REG and the RAND segments were encoded in explicit memory. Together, these data demonstrate an efficient and implicit mechanism that prioritises specific stimulus features to facilitate real-time perception. Listeners anchor to locally repeated motifs of tone sequences (REG segments) when these specific motifs appear across the course of the experiment and do not anchor to RAND segments regardless of their repeated appearance (i.e., in the REP_fixed_ and REP_RAND-fixed_ conditions).

A critical question, then, concerns the type of ‘learning’ this anchoring represents and why the locally repeated motif—the REG segment—in the REP_fixed_ condition drives this anchoring. Does the formation of a stable auditory anchor reflect the slow, cortical consolidation characteristic of long-term statistical learning, or a process of short-term spectro-temporal matching (Agus et al., 2010; Kaernbach, 2004)? We addressed this question by applying inhibitory cTBS over dlPFC—a frontal brain region classically implicated in the active maintenance and manipulation of temporal sequences (Fuster, 2001; Petrides, 2005). If anchoring relies on the dlPFC-dependent consolidation of long-term statistical rules, cTBS should have broken, or at least delayed, the formation of the anchor. It did not. Instead, active disruption of dlPFC not only led to maintenance of the effect, it also significantly accelerated the rate of anchoring in listeners compared to listeners experiencing sham TMS. Whilst aggregate sensitivity (*d’*) was comparable between groups, cTBS facilitated a more rapid transition to a near-zero floor in errors (FAs). This suggests that dlPFC may modulate implicit learning by inhibiting a prefrontal ‘monitor’, and the brain may more efficiently offload fixed auditory information to long-term memory pathways.

Consistent with this interpretation, time-course analysis of the REP_novel_ condition showed that whilst performance in the sham group exhibited a pronounced reduction in performance (detecting identical sequences in a trial), the active cTBS group remained remarkably stable, albeit at an initially higher error rate. This indicates that while the dlPFC is necessary for the initial precision of online structural comparison, its continued activity leads to the accumulation of top-down modulation that eventually causes performance to degrade. Conversely, inhibition of dlPFC appears to protect against any decay in performance over time but trades initial accuracy for stability. Thus, repeated exposure to the anchor (the REG segment of REP_fixed_) effectively offloads the computational burden from prefrontal cortex, allowing implicit, or secondary memory networks to drive stability of the percept without prefrontal interference.

### Perceptual mechanisms supporting auditory integration

Recent models of complex sound perception suggest the auditory system flexibly adjusts its integration time to encode statistically coherent patterns (McDermott & Simoncelli, 2011; McWalter & McDermott, 2018; Teng et al., 2016). Our data suggest that the REG segments in our sequences perform this role of a highly salient, recurring feature set. When a sequence is completely novel (as in the REP_novel_ condition), the auditory system must sample, maintain, and compare these transient local features within working memory, a cognitive process to which dlPFC has been tied (Petrides, 2000; Schneegans et al., 2020).

Nevertheless, our data are also explicable in terms of dlPFC providing the precision required for initial comparison of novel events, but that activity originating in this pathway competes with faster, more efficient implicit pathways during the acquisition of stable anchors. When the REG segment in our paradigm reliably repeats across trials, computational reliance on dlPFC is reduced, and the sequence is consolidated into memory via broader cortical and subcortical networks—likely involving secondary auditory areas, corticofugal efferent pathways, and thalamus—that successfully bypass prefrontal control (Antunes & Malmierca, 2021; Bajo et al., 2019; Barascud et al., 2016). This flexible, dlPFC-dependent weighting mechanism may be particularly critical in naturalistic environments with constantly changing sensory input. Without a reliable cross-trial anchor to guide implicit integration, the brain must continuously deploy prefrontal resources to hold and compare fleeting sensory evidence in real time.

### What stimulus features are being learned in an auditory perceptual anchoring task?

Despite the robustness of the anchoring effect we observed, our reliance on strictly repeating pure-tone sequences potentially limits interpreting these findings as a direct reflection of long-term statistical learning. In the REP_fixed_ condition, sequence structure and statistical probability are perfectly correlated because the sequence is identical across trials, the conditional probability of the next tone is always 1.0. Importantly, though, this holds for both the RAND and the REG segments of the sequences, but only the REG segment—which contains a locally repeating motif—appears to drive anchoring. This suggests the rapid stabilisation we observed reflects a highly efficient, short-term spectro-temporal matching process—driven by local (in time) repetition of the motif in the REG segment—rather than the gradual extraction of underlying probabilistic rules from a varied distribution (Aslin et al., 1998; Saffran et al., 1996). Anchoring in our paradigm seems to require segments not simply to be fixed (multiple exposures over time), nor simply to be repeated (i.e. identical across paired sequences), but to contain a local repetition (i.e., locally repeating motifs in REG segments). RAND segments, which were otherwise equally deterministic as REG segments, did not contain this local repetition. To separate short-term, spectro-temporal feature matching from the longer-term mechanism of statistical learning—if such a separation exists—future anchoring paradigms might benefit from employing more subtle probabilistic manipulations of complex stimuli. Evaluating whether the dlPFC remains disengaged when transitional probabilities are less deterministic will be crucial for mapping the neural boundaries for statistical learning (Bastug et al., 2026; Bonetti et al., 2024; Heilbron & Chait, 2018).

In a recent study exploring listeners’ abilities to adapt to reverberant room acoustics (Hernández-Pérez et al., 2025), we addressed directly the issue of potential abstract acoustic features driving performance in implicit learning tasks. Using a ‘bait and switch’ paradigm in which listeners were required to report words in sentences controlled for semantic content, with room reverberation (defined in terms of the RT_60_) the dependent variable, and talker and phrase length random variables, we found listening performance—quantified in terms of correct words reported—to improve over time, suggesting a benefit of implicitly learning the room reverberation, an abstract stimulus feature defined by its (relatively) long-term statistics. This provides one means of overcoming listeners—ostensibly tasked with learning long-term statistics—nevertheless seeking, and sourcing, short-term spectro-temporal features of sounds to complete a task. This perspective also conforms to phenomena such as auditory pareidolia in which, for example, continuous exposure to white-noise sounds can lead to listeners reporting the experience of structured acoustic patterns, including speech (Nees & Phillips, 2015).

In natural auditory scenes, soundscapes are often fluid, overlapping, and constantly shifting, requiring listeners continuously to adapt to novel information while simultaneously tracking underlying statistical regularities (McDermott et al., 2013). We demonstrated that dlPFC is a significant node in this process, specifically monitoring the short-term computations required when stable structural anchors are absent. Extending this framework to individuals with dyslexia, for example, for whom perceptual anchoring has been demonstrated to be impaired (Ahissar, 2007; Ahissar et al., 2006; Banai & Ahissar, 2010; Daikhin et al., 2017; Shulver & Badcock, 2021), changes in auditory task performance may not strictly reflect a lure to implicitly consolidate long-term rules, but rather some fundamental variation in dlPFC-mediated online maintenance and comparison of novel sensory input. Ultimately, distinguishing between these short-term computational limits and long-term consolidation mechanisms will be essential for refining models of auditory cognition and developing targeted interventions.

### Inclusion and Ethics statement

This study was approved by the Human Research Ethics Committee of Macquarie University (ref: 5201833344987). Each participant signed a written informed consent form and was given financial remuneration for their time.

