## Supplemental Information for "Disruption of prefrontal cortex accelerates sensory anchoring but dissociates online recognition from implicit consolidation of sound sequences"

#### Experiment 1 Training Results

Performance for training was calculated as the overall proportion correct. Training performance showed a reduction in average recognition of $REP_{novel}$ ($M = 0.76$, $SD = 0.14$) compared to $NonREP_{novel}$ ($M = 0.89$, $SD = 0.12$) stimuli, but with an identical range in scores (0.62-1.00). The overall reduction in performance in the $REP_{novel,}$ condition, relative to the $NonREP_{novel}$ condition, suggests that participants were better able to correctly reject non-repeating sequences relative to detecting novel repeating sequences.

#### Experiment 1 Test Results

##### *Sensitivity and Decision Bias*

Sensitivity (*d'*) and decision bias (*c*) were analysed using a Linear Mixed-Effects Model (LMM) with condition ($REP_{novel}$, $REP_{fixed}$) as a fixed effect and participant as a random intercept. The LMM revealed a significant effect of condition on sensitivity ($b = 1.14$, $SE = 0.25$, $t(14) = 4.63$, $p < .001$, with $REP_{fixed}$ ($Mdn = 3.35$, IQR [2.24, 4.38]) showing substantially greater sensitivity than $REP_{novel}$ ($Mdn = 1.88$, IQR [1.43, 2.56]). Decision bias showed a corresponding shift, with $REP_{fixed}$ listeners adopting a more liberal criterion ($Mdn = −0.82$, IQR [−1.10, −0.19]) relative to $REP_{novel}$ ($Mdn = 0.00$, IQR [−0.34, 0.19]). Note that shifts in decision bias are mathematically proportional to shifts in sensitivity within this design, as both conditions share a common $NonREP_{novel}$ false-alarm baseline; a separate model table for criterion is therefore not included. Full LMM output for sensitivity is presented in Table S1.

**Table S1**

*Linear Mixed-Effects Model Fixed Effects for Sensitivity (d'), Experiment 1*

| Parameter | *b* | *SE* | *df* | *t* | *p* |
|---|---|---|---|---|---|
| Intercept | 2.18 | 0.32 | 18.83 | 6.89 | < .001 |
| Condition: $REP_{fixed}$ | 1.14 | 0.25 | 14.00 | 4.63 | < .001 |

*Note.* Random effects: participant variance = 1.05, residual variance = 0.46. $N = 30$ observations, 15 participants. Reference condition = $REP_{novel}$.

##### *Growth Curve Analysis*

Trial-by-trial error probabilities were modelled using a Generalised Linear Mixed-Effects Model (GLMM) with a binomial link function. Orthogonal polynomials up to the second order were used to capture linear (overall rate of improvement) and quadratic (stabilisation) trajectories. The linear time term was significant for the $\text{NonREP}_{\text{novel}}$ baseline ($b = -8.68$, $SE = 3.40$, $z = -2.55$, $p = .011$), reflecting a gradual overall reduction in false alarm rate across the experiment. For $\text{REP}_{\text{fixed}}$, the quadratic interaction term was significant ($b = 70.03$, $SE = 32.63$, $z = 2.15$, $p = .032$), consistent with a rapid initial reduction in errors followed by stabilisation. No significant linear or quadratic effects were observed for $\text{REP}_{\text{novel}}$ (linear: $z = 0.57$, $p = .570$; quadratic: $z = 0.19$, $p = .848$), confirming a flat trajectory in the absence of a recurring pattern. Note that the linear interaction term for $\text{REP}_{\text{fixed}}$ approached but did not reach significance ($b = 94.94$, $SE = 54.42$, $z = 1.74$, $p = .081$), reflecting a trend toward an accelerated initial rate of error reduction for predictable sequences. Full model output is presented in Table S2.

**Table S2**

*Generalised Linear Mixed-Effects Model Fixed Effects for Trial-by-Trial Error Probability (GCA), Experiment 1*

| Parameter | *b* | *SE* | *z* | *p* |
|---|---|---|---|---|
| Intercept | −1.45 | 0.20 | −7.26 | < .001 |
| Linear time | −8.68 | 3.40 | −2.55 | .011 |
| Quadratic time | 0.97 | 3.58 | 0.27 | .786 |
| Condition: $\text{REP}_{\text{novel}}$ | 0.03 | 0.55 | 0.06 | .950 |
| Condition: $\text{REP}_{\text{fixed}}$ | −0.16 | 0.84 | −0.19 | .850 |
| Linear time × $\text{REP}_{\text{novel}}$ | 18.40 | 32.39 | 0.57 | .570 |
| Quadratic time × $\text{REP}_{\text{novel}}$ | 3.93 | 20.49 | 0.19 | .848 |
| Linear time × $\text{REP}_{\text{fixed}}$ | 94.94 | 54.42 | 1.74 | .081 |
| Quadratic time × $\text{REP}_{\text{fixed}}$ | 70.03 | 32.63 | 2.15 | .032 |

*Note.* Random effects: participant variance = 0.40. $N = 1{,}200$ observations, 15 participants. Reference condition = $\text{NonREP}_{\text{novel}}$. Probabilities are modelled on the logit scale. × = interaction.

#### *Estimated Marginal Means at Task Completion*

Estimated marginal means (EMMs) at the final trial are presented in Table S3, reflecting predicted error probabilities upon completion of the full experimental block, back-transformed from the logit scale. EMMs for $\text{NonREP}_{\text{novel}}$ are taken at trial 40; $\text{REP}_{\text{novel}}$ and $\text{REP}_{\text{fixed}}$ EMMs are taken at trial 20, corresponding to the number of trials per condition.

**Table S3**

*Estimated Marginal Means at Final Trial by Condition, Experiment 1*

| Condition | *Estimated Prob.* | *SE* | 95% CI |
|---|---|---|---|
| $\text{NonREP}_{\text{novel}}$ | 0.124 | 0.040 | [0.064, 0.227] |
| $\text{REP}_{\text{novel}}$ | 0.192 | 0.068 | [0.092, 0.358] |
| $\text{REP}_{\text{fixed}}$ | 0.066 | 0.042 | [0.018, 0.214] |

*Note.* Probabilities are back-transformed from the logit scale. $\text{NonREP}_{\text{novel}}$ EMM taken at trial 40; $\text{REP}_{\text{novel}}$ and $\text{REP}_{\text{fixed}}$ EMMs taken at trial 20. Full trial-by-trial EMMs are available from the corresponding author upon request.

#### Experiment 2 Training Results

Performance for the training session was calculated as the overall proportion correct, (i.e., the proportion correct for each condition). Training performance for the shorter-duration sequences showed reduced average recognition of $\text{NonREP}_{\text{novel-short}}$ (M = 0.72, SD = 0.16) relative to $\text{REP}_{\text{novel-short}}$ (M = 0.88, SD = 0.19) stimuli. Similarly, training performance showed reduced average recognition of $\text{NonREP}_{\text{novel}}$ (M = 0.62, SD = 0.23) relative to $\text{REP}_{\text{novel}}$ (M = 0.76, SD = 0.15) stimuli. The overall difference in performance between sequences suggests that the shorter duration sequences were easier to discriminate than longer duration ones, and $\text{REP}_{\text{novel}}$ conditions were easier to discriminate than the $\text{NonREP}_{\text{novel}}$, conditions

### Experiment 2 Test Results

#### *Sensitivity and Decision Bias*

Sensitivity (*d'*) and decision bias (*c*) were analysed using a Linear Mixed-Effects Model (LMM) with condition ($\text{REP}_{\text{novel}}$, $\text{REP}_{\text{fixed}}$, $\text{REP}_{\text{REG-fixed}}$, $\text{REP}_{\text{RAND-fixed}}$) as a fixed effect and participant as a random intercept. The LMM revealed a significant main effect of condition on sensitivity. Relative to the $\text{REP}_{\text{novel}}$ baseline (*Mdn* = 1.60, IQR [1.10, 2.02]), $\text{REP}_{\text{fixed}}$ showed significantly greater sensitivity (*Mdn* = 2.38, IQR [1.94, 3.78]; $b = 1.09$, $SE = 0.41$, $t(39) = 2.69$, $p = .010$, $d = 1.02$), confirming the anchoring effect in this cohort. $\text{REP}_{\text{REG-fixed}}$ showed a profound collapse in sensitivity (*Mdn* = −0.59, IQR [−0.96, 0.14]; $b = -2.05$, $SE = 0.41$, $t(39) = -5.05$, $p < .001$, $d = 1.91$), performing significantly below the $\text{REP}_{\text{novel}}$ baseline. $\text{REP}_{\text{RAND-fixed}}$ did not differ significantly from $\text{REP}_{\text{novel}}$ (*Mdn* = 1.22, IQR [0.87, 1.64]; $b = -0.34$, $SE = 0.41$, $t(39) = -0.83$, $p = .412$, $d = 0.31$). Decision bias mirrored sensitivity: $\text{REP}_{\text{REG-fixed}}$ listeners adopted a pronounced conservative criterion (*Mdn* = 0.92, IQR [0.67, 1.02]), while $\text{REP}_{\text{fixed}}$ shifted to a liberal criterion (*Mdn* = −0.52, IQR [−0.82, −0.29]), and $\text{REP}_{\text{novel}}$ (*Mdn* = −0.04, IQR [−0.47, 0.31]) and $\text{REP}_{\text{RAND-fixed}}$ (*Mdn* = 0.08, IQR [0.02, 0.16]) remained near zero. Note that shifts in decision bias are mathematically proportional to shifts in sensitivity within this design, as conditions share a common $\text{NonREP}_{\text{novel}}$ false-alarm baseline; a separate model table for criterion is therefore not included. Full LMM output is presented in Table S4.

**Table S4**

*Linear Mixed-Effects Model Fixed Effects for Sensitivity (d'), Experiment 2*

| *Parameter* | *b* | *SE* | *df* | *t* | *p* | *d* |
|---|---|---|---|---|---|---|
| Intercept [$\text{REP}_{\text{novel}}$] | 1.71 | 0.33 | 45.19 | 5.26 | < .001 | — |
| Condition: $\text{REP}_{\text{fixed}}$ | 1.09 | 0.41 | 39.00 | 2.69 | .010 | 1.02 |
| Condition: $\text{REP}_{\text{REG-fixed}}$ | −2.05 | 0.41 | 39.00 | −5.05 | < .001 | 1.91 |
| Condition: $\text{REP}_{\text{RAND-fixed}}$ | −0.34 | 0.41 | 39.00 | −0.83 | .412 | 0.31 |

*Note.* Random effects: participant variance = 0.33, residual variance = 1.16. $N = 56$ observations, 14 participants. Reference condition = $\text{REP}_{\text{novel}}$. Cohen's *d* computed as |estimate| / residual SD. Intercept represents $\text{REP}_{\text{novel}}$ mean; *d* not applicable.

***Growth Curve Analysis***

Trial-by-trial error probabilities were modelled using a GLMM with a binomial link function and orthogonal polynomials up to second order. Significant effects emerged for $REP_{fixed}$: the linear interaction term indicated an accelerated rate of error reduction ($b = 963.59$, $SE = 425.55$, $z = 2.26$, $p = .024$), and the quadratic interaction term confirmed subsequent stabilisation ($b = 342.44$, $SE = 154.52$, $z = 2.22$, $p = .027$). The condition main effect for $REP_{fixed}$ was also significant ($b = 10.52$, $SE = 5.26$, $z = 2.00$, $p = .046$), reflecting an overall elevation in the intercept for predictable sequences. No significant linear or quadratic effects were observed for $REP_{novel}$, $REP_{REG\text{-}fixed}$, or $REP_{RAND\text{-}fixed}$ (all *p*s > .30), confirming flat trajectories for these conditions. Full model output is presented in Table S5.

**Table S5**

*Generalised Linear Mixed-Effects Model Fixed Effects for Trial-by-Trial Error Probability (GCA), Experiment 2*

| Parameter | *b* | *SE* | *z* | *p* |
|---|---|---|---|---|
| Intercept | −1.12 | 0.09 | −12.35 | < .001 |
| Linear time | −0.06 | 3.39 | −0.02 | .986 |
| Quadratic time | 4.25 | 2.91 | 1.46 | .145 |
| Condition: $REP_{novel}$ | −2.61 | 3.37 | −0.77 | .439 |
| Condition: $REP_{fixed}$ | 10.52 | 5.26 | 2.00 | .046 |
| Condition: $REP_{REG\text{-}fixed}$ | 1.41 | 3.46 | 0.41 | .684 |
| Condition: $REP_{RAND\text{-}fixed}$ | 2.40 | 3.01 | 0.80 | .425 |
| Linear time × $REP_{novel}$ | −211.87 | 266.77 | −0.79 | .427 |
| Quadratic time × $REP_{novel}$ | −95.85 | 97.62 | −0.98 | .326 |
| Linear time × $REP_{fixed}$ | 963.59 | 425.55 | 2.26 | .024 |
| Quadratic time × $REP_{fixed}$ | 342.44 | 154.52 | 2.22 | .027 |

| Parameter | *b* | *SE* | *z* | *p* |
|---|---|---|---|---|
| Linear time × $REP_{REG\text{-}fixed}$ | 129.62 | 274.34 | 0.47 | .637 |
| Quadratic time × $REP_{REG\text{-}fixed}$ | 47.32 | 98.98 | 0.48 | .633 |
| Linear time × $REP_{RAND\text{-}fixed}$ | 50.66 | 238.18 | 0.21 | .832 |
| Quadratic time × $REP_{RAND\text{-}fixed}$ | 16.39 | 86.28 | 0.19 | .849 |

*Note.* Random effects: participant variance = 0.008. $N$ = 2,240 observations, 14 participants. Reference condition = $NonREP_{novel}$. × = interaction.

#### ***Estimated Marginal Means at Task Completion***

Estimated marginal means (EMMs) at the final trial are presented in Table S6, reflecting predicted error probabilities upon completion of the full experimental block, back-transformed from the logit scale. EMMs for $NonREP_{novel}$ are taken at trial 80; all REP condition EMMs are taken at trial 20, corresponding to the number of trials per condition.

**Table S6**

*Estimated Marginal Means at Final Trial by Condition, Experiment 2*

| Condition | *Estimated Prob.* | *SE* | 95% CI |
|---|---|---|---|
| $NonREP_{novel}$ | 0.298 | 0.041 | [0.224, 0.385] |
| $REP_{novel}$ | 0.266 | 0.072 | [0.149, 0.427] |
| $REP_{fixed}$ | 0.158 | 0.068 | [0.064, 0.339] |
| $REP_{REG\text{-}fixed}$ | 0.216 | 0.068 | [0.112, 0.376] |
| $REP_{RAND\text{-}fixed}$ | 0.658 | 0.077 | [0.495, 0.790] |

*Note.* Probabilities are back-transformed from the logit scale. $NonREP_{novel}$ EMM taken at trial 80; all REP condition EMMs taken at trial 20. Full trial-by-trial EMMs are available from the corresponding author upon request.

**Experiment 3 Training Results**

Performance for training was calculated as the overall proportion correct. Training performance for the shorter duration noises showed a reduction in average recognition of $\text{NonREP}_{\text{novel-short}}$ (M = 0.76, SD = 0.21) relative to $\text{REP}_{\text{novel-short}}$ (M = 0.76, SD = 0.20) stimuli (Figure 4.3. Panel A). Similarly, training performance showed a reduction in average recognition of $\text{NonREP}_{\text{novel}}$ (M = 0.86, SD = 0.12) relative to $\text{REP}_{\text{novel}}$ (M = 0.69, SD = 0.23) stimuli (see Figure 4.3 Panel A). The overall difference in performance between noises suggests that the shorter duration noises were easier to discriminate than longer duration noises, and $\text{REP}_{\text{novel}}$ conditions were easier to discriminate than the $\text{NonREP}_{\text{novel}}$, conditions.

**Experiment 3 Test Results**

***Sensitivity and Decision Bias***

Sensitivity (*d'*) was analysed using a Linear Mixed-Effects Model (LMM) with condition ($\text{REP}_{\text{novel}}$, $\text{REP}_{\text{fixed}}$) and group (sham, active TMS) as fixed effects, their interaction, and participant as a random intercept. The LMM revealed a significant main effect of condition: listeners showed substantially greater sensitivity for $\text{REP}_{\text{fixed}}$ relative to $\text{REP}_{\text{novel}}$ ($b = 1.21$, $SE = 0.31$, $t(18) = 3.84$, $p = .001$, $d = 1.72$). The main effect of group was not significant ($b = -0.003$, $SE = 0.38$, $t(32.59) = -0.007$, $p = .995$, $d < .01$), nor was the condition × group interaction ($b = 0.04$, $SE = 0.44$, $t(18) = 0.09$, $p = .933$, $d = 0.05$), confirming that overall anchoring proficiency was preserved following cTBS. Tukey-corrected pairwise contrasts are reported in Table S8. Note: descriptive statistics for *d'* and *c* by condition and group are presented in Table S7. Full LMM output is presented in Table S8.

**Table S7**

*Descriptive Statistics for Sensitivity (d') and Decision Bias (c) by Condition and Group, Experiment 3*

| Condition | Group | *d'* | | | *c* | | |
|---|---|---|---|---|---|---|---|
| | | *Mdn* | *Q1* | *Q3* | *Mdn* | *Q1* | *Q3* |
| $\text{REP}_{\text{novel}}$ | Sham | 1.56 | 1.32 | 1.80 | 0.238 | −0.165 | 0.372 |
| $\text{REP}_{\text{novel}}$ | TMS | 1.46 | 1.05 | 2.10 | 0.124 | 0.060 | 0.337 |
| $\text{REP}_{\text{fixed}}$ | Sham | 2.66 | 2.44 | 3.54 | −0.317 | −1.260 | −0.019 |

| Condition | Group | $d'$ | | | $c$ | | |
|---|---|---|---|---|---|---|---|
| | | *Mdn* | *Q1* | *Q3* | *Mdn* | *Q1* | *Q3* |
| $\text{REP}_{\text{fixed}}$ | TMS | 2.64 | 2.38 | 3.32 | −0.302 | −0.578 | −0.254 |

*Note. Mdn* = median; IQR = interquartile range. $d'$ = sensitivity; $c$ = decision bias (criterion). Values derived from R output.

**Table S8**

*Linear Mixed-Effects Model Fixed Effects for Sensitivity (d'), Experiment 3*

| Parameter | *b* | *SE* | *df* | *t* | *p* | *d* |
|---|---|---|---|---|---|---|
| Intercept [$\text{REP}_{\text{novel}}$ sham] | 1.58 | 0.27 | 32.59 | 5.84 | < .001 | — |
| Condition: $\text{REP}_{\text{fixed}}$ | 1.21 | 0.31 | 18.00 | 3.84 | .001 | 1.72 |
| Group: TMS | −0.003 | 0.38 | 32.59 | −0.007 | .995 | < .01 |
| Condition: $\text{REP}_{\text{fixed}}$ × Group: TMS | 0.04 | 0.44 | 18.00 | 0.09 | .933 | 0.05 |

*Note.* Random effects: participant variance = 0.24, residual variance = 0.49. $N = 40$ observations, 20 participants (10 sham, 10 active TMS). Reference condition = $\text{REP}_{\text{novel}}$ sham. Cohen's $d$ computed as |estimate| / residual SD. Intercept represents $\text{REP}_{\text{novel}}$ sham mean; $d$ not applicable.

**Table S9**

*Tukey-Corrected Pairwise Contrasts for Sensitivity (d') by Condition and Group, Experiment 3*

| Contrast | Estimate | *SE* | *df* | *t* | *p* | *d* |
|---|---|---|---|---|---|---|
| REP(novel) sham − REP(fixed) sham | −1.21 | 0.31 | 18.0 | −3.84 | .006 | 1.72 |
| REP(novel) sham − REP(novel) TMS | 0.003 | 0.38 | 32.6 | 0.01 | 1.000 | < .01 |

| Contrast | Estimate | *SE* | *df* | *t* | *p* | *d* |
|---|---|---|---|---|---|---|
| REP(novel) sham − REP(fixed) TMS | −1.24 | 0.38 | 32.6 | −3.25 | .014 | 1.77 |
| REP(fixed) sham − REP(novel) TMS | 1.21 | 0.38 | 32.6 | 3.17 | .017 | 1.72 |
| REP(fixed) sham − REP(fixed) TMS | −0.04 | 0.38 | 32.6 | −0.09 | 1.000 | 0.05 |
| REP(novel) TMS − REP(fixed) TMS | −1.25 | 0.31 | 18.0 | −3.96 | .005 | 1.77 |

*Note. p* values adjusted using Tukey method for a family of 4 estimates. Degrees of freedom estimated via Kenward-Roger method. Cohen's *d* computed as |estimate| / residual SD (residual SD = 0.70).

#### *Growth Curve Analysis*

Four GLMM analyses were conducted to characterise learning dynamics. First, a full model including condition, group, time (orthogonal polynomials to second order), and all interactions was fitted across all trials. Second, a model restricted to early trials (1–10) was fitted to examine transient effects. Third, a focused model on $REP_{novel}$ data only was fitted to characterise within-condition divergence between groups. Full model outputs are presented in Tables S9–S11 respectively.

**Table S10**

*Generalised Linear Mixed-Effects Model Fixed Effects for Trial-by-Trial Error Probability (Full GCA Model), Experiment 3*

| Parameter | *b* | *SE* | *z* | *p* |
|---|---|---|---|---|
| Intercept | −1.33 | 0.17 | −7.76 | < .001 |
| Linear time | −0.63 | 4.55 | −0.14 | .890 |
| Quadratic time | −3.90 | 4.61 | −0.85 | .398 |

| Parameter | *b* | *SE* | *z* | *p* |
|---|---|---|---|---|
| Condition: $REP_{novel}$ | 0.40 | 0.27 | 1.47 | .142 |
| Condition: $REP_{fixed}$ | −0.12 | 0.31 | −0.39 | .700 |
| Group: TMS | −0.20 | 0.24 | −0.82 | .415 |
| Linear time × $REP_{novel}$ | 17.99 | 22.63 | 0.80 | .427 |
| Quadratic time × $REP_{novel}$ | −4.40 | 15.39 | −0.29 | .775 |
| Linear time × $REP_{fixed}$ | 96.22 | 24.79 | 3.88 | < .001 |
| Quadratic time × $REP_{fixed}$ | 65.18 | 17.21 | 3.79 | < .001 |
| Linear time × Group: TMS | 0.46 | 6.56 | 0.07 | .944 |
| Quadratic time × Group: TMS | 5.40 | 6.62 | 0.82 | .415 |
| Condition: $REP_{novel}$ × Group: TMS | 0.26 | 0.36 | 0.71 | .480 |
| Condition: $REP_{fixed}$ × Group: TMS | −1.07 | 0.53 | −2.04 | .041 |
| Linear time × $REP_{novel}$ × Group: TMS | −8.24 | 29.62 | −0.28 | .781 |
| Quadratic time × $REP_{novel}$ × Group: TMS | 14.05 | 19.67 | 0.71 | .475 |
| Linear time × $REP_{fixed}$ × Group: TMS | −94.60 | 42.47 | −2.23 | .026 |
| Quadratic time × $REP_{fixed}$ × Group: TMS | −34.79 | 26.62 | −1.31 | .191 |

*Note.* Random effects: participant variance = 0.20. *N* = 3,200 observations, 20 participants. Reference condition = $NonREP_{novel}$ sham. × = interaction.

**Table S11**

*Generalised Linear Mixed-Effects Model Fixed Effects for Early-Phase Trial-by-Trial Error Probability (Trials 1–10, Full Model), Experiment 3*

| Parameter | *b* | *SE* | *z* | *p* |
|---|---|---|---|---|
| Intercept | −1.07 | 0.55 | −1.93 | .053 |
| Trial | −0.05 | 0.09 | −0.61 | .543 |
| Condition: $REP_{novel}$ | −1.60 | 0.83 | −1.93 | .054 |
| Condition: $REP_{fixed}$ | 0.17 | 0.84 | 0.20 | .843 |
| Group: TMS | −0.86 | 0.82 | −1.05 | .293 |
| Trial × Condition: $REP_{novel}$ | 0.27 | 0.13 | 2.15 | .032 |
| Trial × Condition: $REP_{fixed}$ | −0.29 | 0.18 | −1.66 | .096 |
| Trial × Group: TMS | 0.15 | 0.12 | 1.23 | .219 |
| Condition: $REP_{novel}$ × Group: TMS | 2.75 | 1.12 | 2.45 | .014 |
| Condition: $REP_{fixed}$ × Group: TMS | 0.29 | 1.21 | 0.24 | .812 |
| Trial × $REP_{novel}$ × Group: TMS | −0.39 | 0.17 | −2.24 | .025 |
| Trial × $REP_{fixed}$ × Group: TMS | 0.02 | 0.23 | 0.09 | .931 |

*Note.* Random effects: participant variance = 0.37. *N* = 600 observations, 20 participants. Reference condition = $NonREP_{novel}$ sham. × = interaction.

**Table S12**

*Generalised Linear Mixed-Effects Model Fixed Effects for REPnovel Early-Phase Error Probability (Trials 1–10), Experiment 3*

| Parameter | *b* | *SE* | *z* | *p* |
|---|---|---|---|---|
| Intercept | −2.85 | 0.74 | −3.85 | < .001 |
| Trial | 0.23 | 0.10 | 2.42 | .016 |
| Group: TMS | 2.01 | 0.93 | 2.17 | .030 |

| Parameter | *b* | *SE* | *z* | *p* |
|---|---|---|---|---|
| Trial × Group: TMS | −0.25 | 0.13 | −1.99 | .047 |

*Note.* Random effects: participant variance = 0.77. *N* = 200 observations, 20 participants ($REP_{novel}$ trials only). Reference group = sham. Positive trial slope for sham group indicates rising error rate across early trials; significant trial × group interaction indicates divergent trajectories between groups.

#### ***Estimated Marginal Means at Task Completion***

Estimated marginal means (EMMs) at the final trial are presented in Table S12, reflecting predicted error probabilities back-transformed from the logit scale, separated by condition and group. EMMs for $NonREP_{novel}$ are taken at trial 80; REP condition EMMs are taken at trial 40, corresponding to the number of trials per condition per group.

**Table S13**

*Estimated Marginal Means at Final Trial by Condition and Group, Experiment 3*

| Condition | Group | *Estimated Prob.* | *SE* | 95% CI |
|---|---|---|---|---|
| $NonREP_{novel}$ | Sham | 0.175 | 0.043 | [0.106, 0.274] |
| $REP_{novel}$ | Sham | 0.346 | 0.060 | [0.239, 0.471] |
| $REP_{fixed}$ | Sham | 0.143 | 0.041 | [0.080, 0.242] |
| $NonREP_{novel}$ | TMS | 0.189 | 0.046 | [0.115, 0.295] |
| $REP_{novel}$ | TMS | 0.269 | 0.060 | [0.169, 0.401] |
| $REP_{fixed}$ | TMS | 0.036 | 0.018 | [0.013, 0.093] |

*Note.* Probabilities are back-transformed from the logit scale. $NonREP_{novel}$ EMMs taken at trial 80; REP condition EMMs taken at trial 40. Full trial-by-trial EMMs are available from the corresponding author upon request.

**Code and Data Availability**

Analysis code and any additional analyses not reported in the manuscript or supplemental material are available from the corresponding author upon reasonable request. Requests should be directed to Kurt D Shulver.
